# The C-terminal domain of Hsp70 is responsible for paralog-specific regulation of ribonucleotide reductase

**DOI:** 10.1101/2022.02.08.479504

**Authors:** Laura E. Knighton, Nitika, Siddhi Omkar, Andrew. W. Truman

## Abstract

The Hsp70 family of molecular chaperones is well-conserved and expressed in all organisms. In budding yeast, cells express four highly similar cytosolic Hsp70s Ssa1, 2, 3 and 4 which arose from gene duplication. Ssa1 and 2 are constitutively expressed while Ssa3 and 4 are induced upon heat shock. Recent evidence suggests that despite their amino acid similarity, these Ssas have unique roles in the cell. Here we examine the relative importance of Ssa1-4 in the regulation of the enzyme ribonucleotide reductase (RNR). We demonstrate that cells expressing either Ssa3 or Ssa4 as their sole Ssa are compromised for their resistance to DNA damaging agents and activation of DDR transcription. In addition, we show that the steady state levels and stability of RNR small subunits Rnr2 and Rnr4 are reduced in Ssa3 or Ssa4-expressing cells, a result of decreased Ssa-RNR interaction. Interaction between the Hsp70 co-chaperone Ydj1 and RNR is correspondingly decreased in cells only expressing Ssa3 and 4. Through studies of Ssa2/4 domain swap chimeras, we determined that the C-terminal domain of Ssas are the source of this functional specificity. Taking together, our work suggests a distinct role for Ssa paralogs in regulating DNA replication mediated by C-terminus sequence variation.

**Author Summary:** Cells require molecular chaperones to fold proteins into their active conformation. A major mystery however is why cells express so many highly-related and apparently redundant Hsp70 paralogs. We examined the role of four Hsp70 paralogs in budding yeast (Ssa1, 2, 3 and 4) on the activity of the ribonucleotide reductase (RNR complex). Importantly, we demonstrate there is selectivity of RNR subunits for Ssa1 and Ssa2 subunits, which is dictated by the co-chaperone Ydj1. Taken together, our work provides new insight into the functional specificity of Hsp70 paralogs using a native client protein.

## Introduction

Cells in all organisms must be able to cope with stressors that trigger both protein unfolding and misfolding. The core machinery induced in response to challenges of proteostasis are the molecular chaperones or Heat Shock Proteins [1, 2]. The essential chaperone Heat Shock Protein 70 (Hsp70) is a main player in proteostasis, binding nascent chains and responsible for stabilization, folding and degradation of a large majority of the proteome [1, 3, 4]. Structurally, Hsp70 chaperones comprise of three major functional domains; an N-terminal ATPase domain (NBD) connected by a flexible linker to a substrate binding domain (SBD) and a C-terminal domain (CTD) [2]. Binding and hydrolysis of ATP by the N-terminus promotes a range of conformational changes which are transduced through the linker into the C-terminus resulting in clamping of the CTD over the SBD trapping clients and promoting protein folding [2]. Hsp70 requires the assistance of co-chaperone “helper” proteins, comprised of J-proteins and nucleotide exchange factors (NEFs) that facilitate the stimulation of Hsp70 activity and folding of client proteins [3, 5, 6]. Although the exact mechanism by which Hsp70 folds clients remains unclear, recent evidence suggests that Hsp70 works like a molecular “hair straightener”, pulling out kinks in non-optimal protein conformations, allowing correct folding to occur [7].

Given their role in protein folding, it is unsurprising that Hsp70 appears to exist in most organisms studied so far and is essential for their viability [2]. However, the cellular rationale for the large number of highly related paralogs of Hsp70 remains unclear. Budding yeast (*S. cerevisiae*) expresses 14 different Hsp70 paralogs, five of which are localized to specific organelles [8]. Out of the remaining 9 cytosolic Hsp70s, 4 are from the Stress Seventy sub-family A (SSA) comprised of Ssa1, 2, 3 and 4 [8–12]. Previously, Hsp70 paralogs were thought to be functionally indistinguishable apart from spatiotemporal expression patterns, however recent findings suggest unique functions for Ssa paralogs [8, 9].

Ssa paralogs arose from genome duplication and are highly conserved, with Ssa1 sharing 99%, 84% and 85% amino acid identity with Ssa2, 3 and 4, respectively [8]. The most prominent difference between the Ssa1-4 paralogs is their expression levels; Ssa1/2 are expressed constitutively at high levels whereas Ssa3/4 are only expressed during cell stress [8, 13–17]. Although yeast can survive on the loss of any of 3 Ssas if the 4^th^ is expressed at high levels, the phenotypes of these cells vary in terms of heat resistance longevity, ability to fold certain clients [18–21].

In this study, in order to better understand functional differences between Ssa1-4, we utilized the model chaperone client Ribonucleotide Reductase (RNR). RNR is an enzyme that is important for the production of deoxyribonucleotides (dNTPs) which are used in DNA synthesis and repair [22]. RNR is comprised of two diverse subunits, the large subunit R1 (R1 in vertebrates, Rnr1/Rnr3 in yeast) which contains the allosteric regulatory sites [23] and the small subunit R2 (R2/R2B in vertebrates, Rnr2/Rnr4 in yeast) which consists of a cell cycle regulated binuclear iron center and a tyrosyl free radical [24–28]. Due to its crucial role in the maintenance of genome integrity and subsequently cell survival, RNR remains an attractive target in cancer treatments [25, 27, 29]. Several RNR inhibitors have been developed and used in a clinical setting including hydroxyurea (HU), triapine and gemcitabine [25, 30–32].

Previous studies in both yeast and mammalian cells have identified Hsp70 as an important regulator of RNR with small molecule chaperone inhibitors such as 17-AAG promoting RNR subunit degradation [33–35]. Consequently, Hsp70 inhibition sensitizes cancer cells to gemcitabine, which has the potential to be a novel anti-cancer therapeutic [33–38]. Recently, the Hsp70 co-chaperone Ydj1/DNAJA1 (yeast/mammalian) was identified to assist Hsp70 in the regulation of RNR. Lack of Ydj1 in *S. cerevisiae* results in reduced Rnr2 subunit expression and stability [34, 35]. This interaction was found to be conserved in humans, where DNAJA1 and R2B assist in RNR complex stability and activity in mammalian cells [34, 35]. Additionally, inhibition of DNAJA1 with 116-9e, a small molecule inhibitor that blocks Hsp40 binding to Hsp70 through the J-domain resulted in disruption of R2B-DNAJA1 interaction and sensitized cells to HU and triapine [34, 35].

Here we characterize the relative roles of Ssa1, 2, 3, and 4 in regulating ribonucleotide reductase in yeast. We reveal that yeast expressing single Ssas as their sole cytosolic Hsp70 on identical promoters display differing abilities to respond to DNA damaging agents. This can be explained by the loss of RNR subunit stability, occurring due to decreased Ssa-RNR interaction. Finally, we provide evidence that the C-terminal domain of Hsp70 is responsible for selectivity in activating RNR.

## Results

### Ssa paralogs contribute differentially to the resistance to DNA-perturbing agents and the transcriptional response to DNA damage

Previous studies have demonstrated a critical role for Hsp70 and Hsp90 in supporting RNR activity in yeast and mammalian cells [33–35]. In order to dissect the unique roles of the yeast Hsp70 paralogs Ssa1-4 in supporting the DNA damage response (DDR), we screened cells expressing either Ssa1, 2, 3 or 4 (under the constitutive Ssa2 promoter) as the sole cytosolic Hsp70 for growth against various DNA damaging agents including hydroxyurea (HU), 5-fluorouracil (5-FU), hydrogen peroxide (H_2_O_2_) and methyl methanesulfonate (MMS) (Fig. 1A-D). Yeast expressing Ssa1 or Ssa2 were markedly more resistant to all DNA damaging agents compared to Ssa3 or Ssa4 cells (Fig. 1A). Cells expressing Ssa3 or Ssa4 as their sole Ssa displayed an increased sensitivity to HU, 5-FU and H_2_O_2_ but not MMS (Fig. 1A-D). To determine whether this phenotypic difference was due to altered DNA damage response-regulated transcription, we compared induction of *RNR3* in HU-treated Ssa1, 2, 3 and 4 cells (Fig. 1E). Consistent with the phenotypes in Fig. 1, Ssa3 and Ssa4 cells were unable to fully activate DDR transcription, displaying a significant decrease in HU-mediated *RNR3* expression (Fig. 1E).

**Figure 1.**
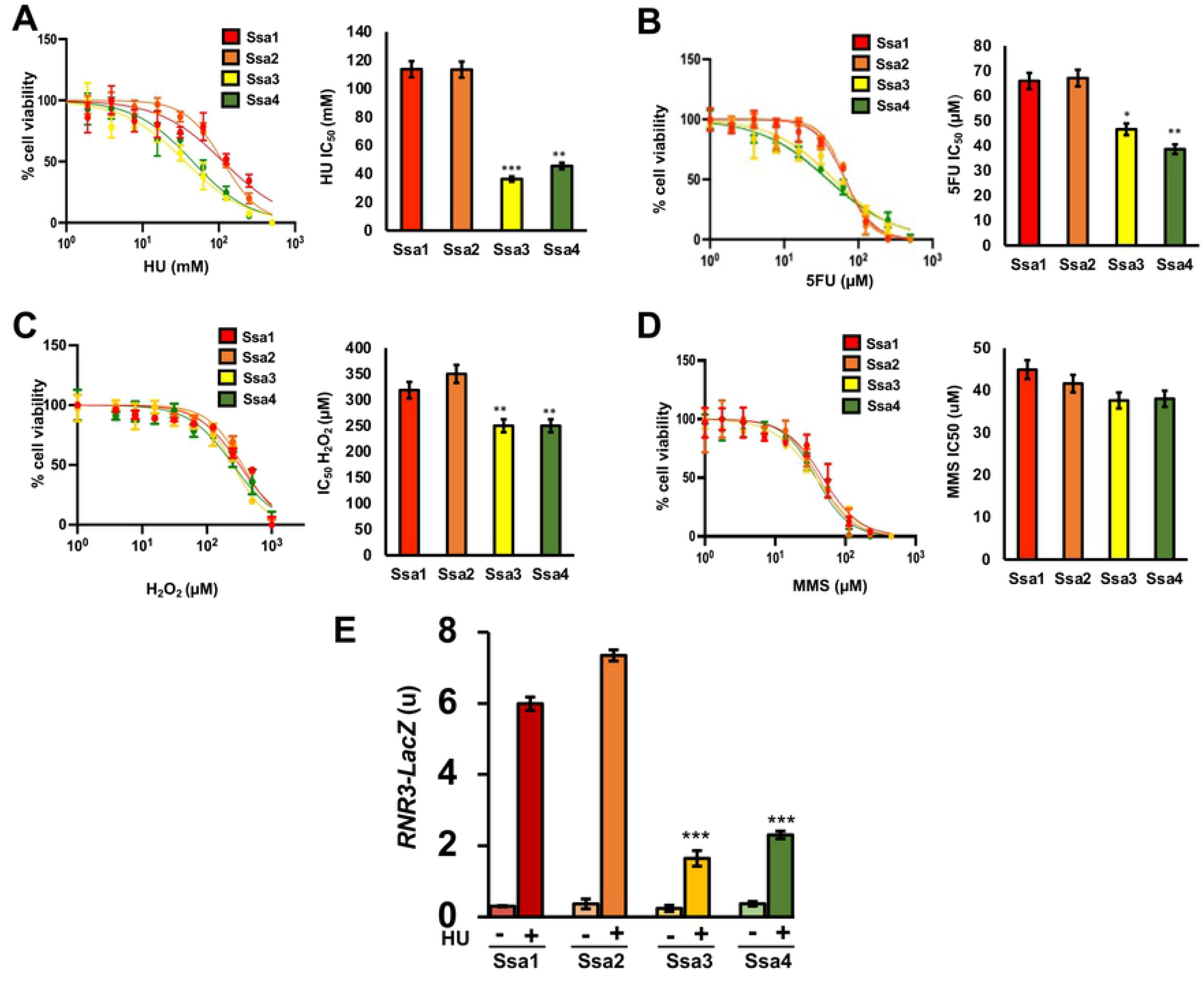
Ssa paralogs confer differential resistance to DNA-damaging agents. (A-D) mid-log *ssa1-4Δ* cells expressing either Ssa1, 2, 3 or 4 were treated with a serial dilution of the indicated drug in a 96-well format for 18 hrs. at which point yeast growth (OD600) was measured via plate reader. Each value represents the mean and standard deviation (error bar) from three independent transformants. Statistical significance were calculated via ANOVA. *, P≤0.05; **, P≤0.01; ***, P≤0.001 as compared to the Ssa2 strain. (E) An *RNR3-LacZ* reporter plasmid was transformed into the indicated yeast strains. Transformants were grown and subjected to either 0 or 200mM HU for 3 hours. β-Galactosidase activity was measured in crude extracts. β-Galactosidase specific activity [-Gal Sp. Act. (U)] is shown on the y axis. Each value represents the mean and standard deviation (error bar) from three independent transformants; *, P≤0.05**, P≤0.01; ***, P≤0.001 as compared to the Ssa2 strain.

### Ribonucleotide reductase subunit levels are compromised in cells solely expressing either Ssa3 or Ssa4

Our previous studies described a role for Hsp70 function in maintaining an active RNR complex in yeast and human cells. To determine whether the inability of Ssa3/4 cells to grow in the presence of DNA-damaging agents could be explained by loss of RNR function, we queried the steady-state levels of Rnr1, Rnr2 and Rnr4 protein in cells expressing single Ssa paralogs. Although Rnr1 levels remained independent of Ssa1 paralog (Fig. 2A), clear differences in Rnr2 and Rnr4 levels were observed. Rnr2 levels in Ssa3/4 cells were significantly lowered in both untreated and treated conditions compared to cells whose primary Hsp70 was Ssa1 or Ssa2 (Fig. 2B). In contrast, while Rnr4 levels in untreated conditions were independent of Ssa version, the levels of HU-induced Rnr4 were substantially decreased in Ssa3 or Ssa4 cells (Fig. 2C).

**Figure 2.**
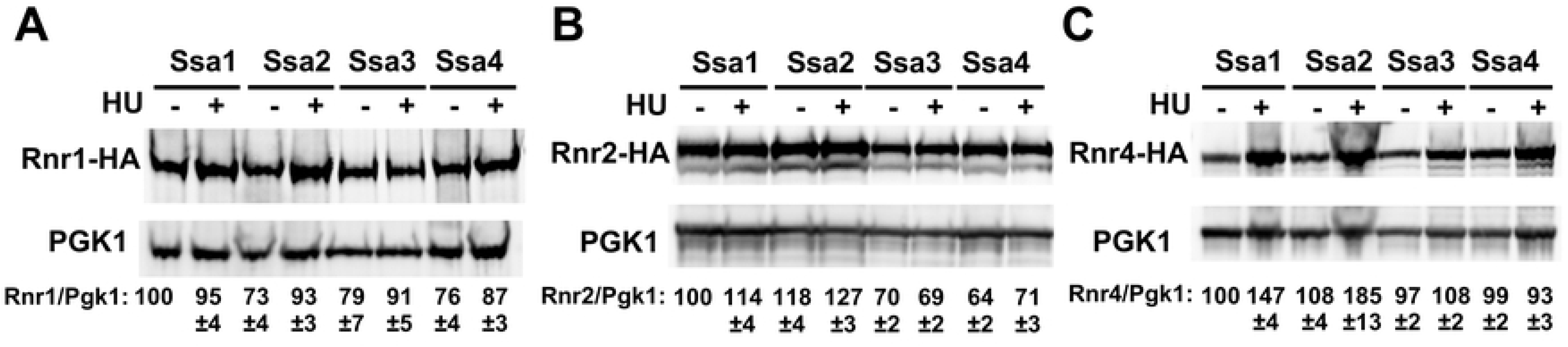
Steady-state levels of RNR small subunits are dependent on Ssa paralogs. *ssa1-4Δ* cells expressing either Ssa1, 2, 3 or 4 and endogenously tagged Rnr1-HA, Rnr2-HA or Rnr4-HA were grown to exponential phase and were either left untreated or were treated with 200mM HU for 3 hours. Cell extracts were obtained, resolved on SDS-PAGE gels and analyzed by immunoblotting with anti-HA and PGK1 antibodies. The ratio of RNR subunit/PGK was quantified and determined from three replicate experiments.

The steady state level of proteins are carefully balanced by both rate of transcription and protein degradation. To determine whether the altered RNR subunit expression observed in Ssa3 and Ssa4 cells was a result of altered transcription, we quantified *RNR1*, *RNR2* and *RNR4* mRNA expression in Ssa-paralog specific yeast using real-time quantitative polymerase chain reaction (RT-qPCR). *RNR1 and RNR2* transcription was independent of Ssa paralog, whereas HU-induced *RNR4* induction was compromised in cells expressing only Ssa3 and Ssa4 (Fig. 3A). To determine whether the protein stability of Rnr1, Rnr2 and Rn4 had also been compromised in Ssa3 and Ssa4-expressing cells, we examined the half-life of RNR subunits by transcriptional shut-off experiments. While Rnr1 stability was independent of Ssa paralog, Rnr2 stability was substantially lowered in Ssa3/4 cells (Fig. 3B). In contrast, Rnr4 stability was only decreased in Ssa3 cells (Fig. 3C).

**Figure 3.**
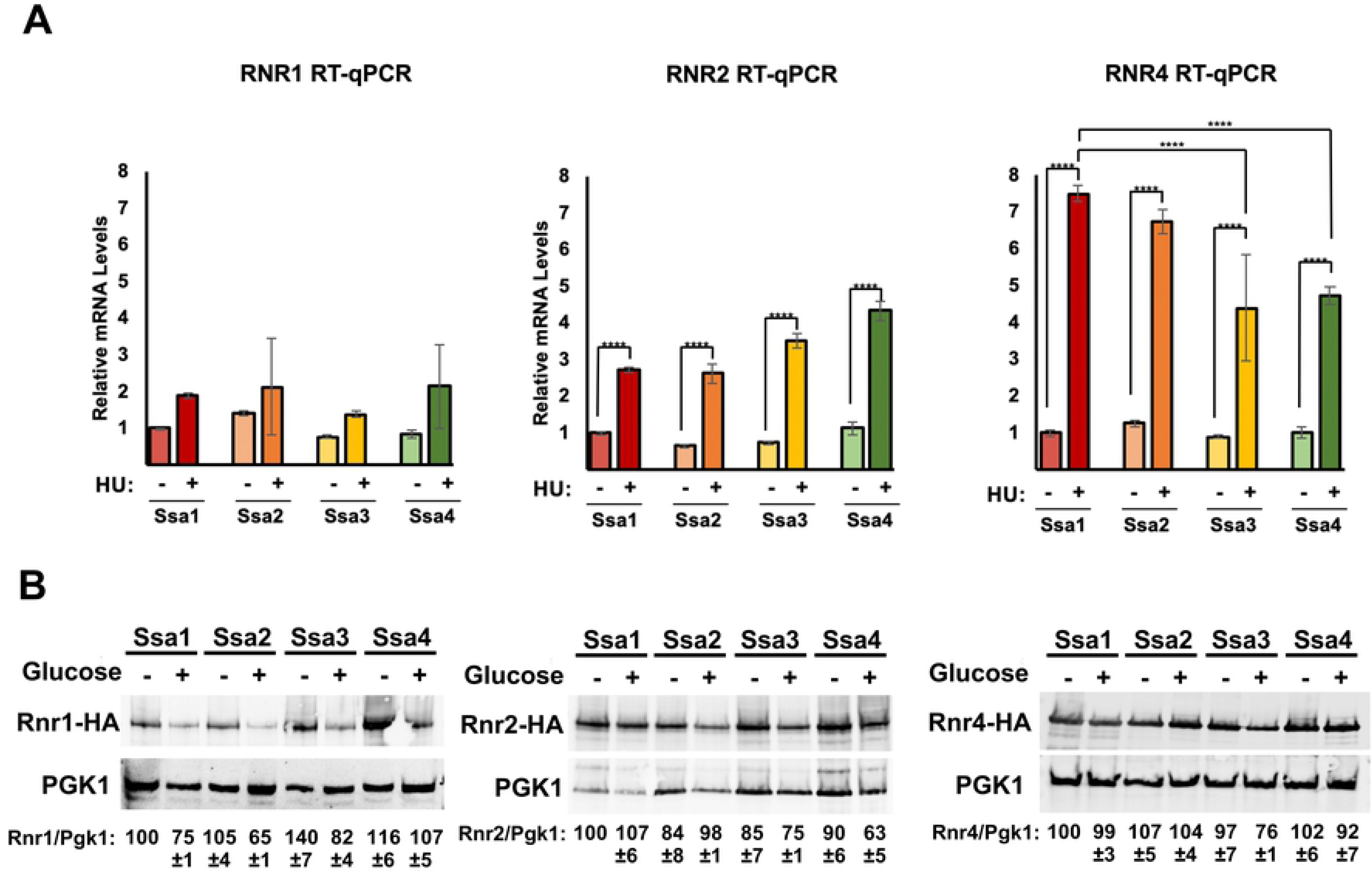
Transcription and stability of RNR subunits are altered in cells only expressing one Ssa paralog. (A) Quantitation of *RNR1*, *RNR2* and *RNR4* mRNA expression in Ssa1, 2,3 or 4-expressing yeast. Levels of *RNR1*, *RNR2* and *RNR4* mRNAs in Ssa1, 2,3 or 4 cells were determined by reverse transcription and RT-qPCR. Signals of *RNR1*, *RNR2* and *RNR4* were normalized against that of *ACT1* in each strain, and the resulting ratios in Ssa1 cells were arbitrarily defined as onefold. Data are the average and SD from three replicates *, P≤0.05**, P≤0.01; ***, P≤0.001 as compared to indicated strains. (B) RNR subunit stability in yeast expressing single Ssa paralogs. Ssa1, 2,3 or 4-expressing cells transformed with either pGAL1-HA-Rnr1, 2 or 4 plasmids were grown to mid-log phase in YP Galactose medium. RNR expression was shut off by addition of 2% glucose to cultures. Cell lysates from these samples were analyzed over time by Western Blotting for the stability of HA-RNR subunit (HA antibody) and loading control (PGK1). The ratio of RNR/PGK was quantified and determined from three replicate experiments.

### RNR subunits display a binding preference for Ssa1 and Ssa2

Hsp90, Hsp70 and Hsp40 proteins in yeast and mammalian cells bind RNR subunit components [33–35]. In order to determine if the observed RNR instability in Ssa3/4 cells was a consequence of decreased RNR-chaperone interaction, we assessed the physical interaction of Rnr1, Rnr2 and Rnr4 proteins with Ssa1, 2, 3, and 4 by yeast two-hybrid analysis. In both experiments, Rnr1 interacted equally with all four Ssas (Fig. 4A), whereas both Rnr2 and Rnr4 displayed a clear binding preference for Ssa1 and Ssa2 (Fig. 4B and 4C).

**Figure 4.**
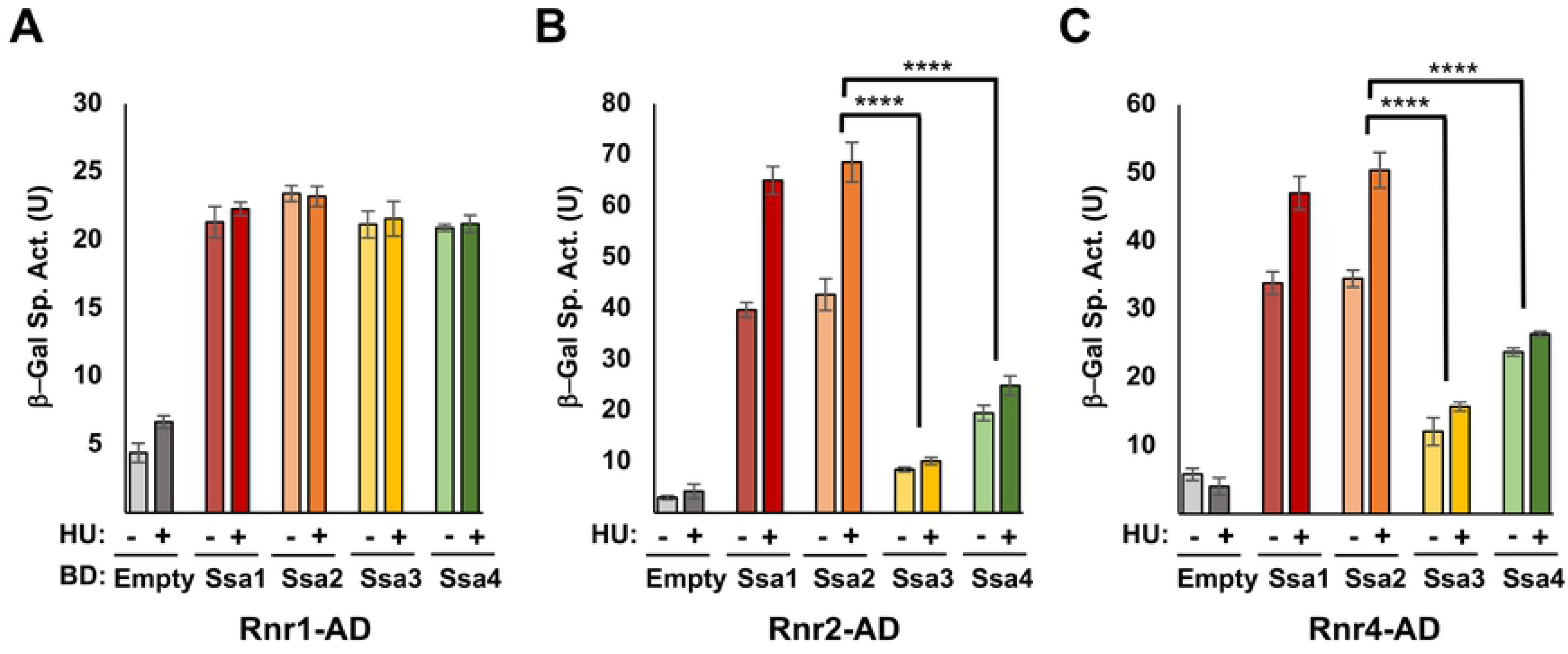
RNR small subunits display binding selectivity for Ssa paralogs. PJ694a/ α cells were transformed with transformed with the appropriate AD-RNR and BD-Ssa fusions. Cells were grown in selective media to the mid-log phase at which point protein was extracted and analyzed for β-galactosidase activity. Each value represents the mean and standard deviation (error bar) from three independent transformants. Statistical significance between samples was calculated as above. * P≤0.05**, P≤0.01; ***, P≤0.001 compared to indicated strains.

### RNR-Ydj1 interaction is weaker in cells solely expressing Ssa3 or Ssa4

Recent studies have revealed that Hsp70 paralogs display differing affinities for their associated co-chaperones [21, 39]. In light of our previous work indicating that Ydj1 directly regulates RNR activity [34], we sought to determine whether Ydj1 association with RNR was reduced in Ssa3/4-expressing cells. We immunoprecipitated HA-tagged Rnr1, Rnr2 or Rnr4 from Ssa1, Ssa2, Ssa3 or Ssa4 cells and assessed the association of Ydj1 via Western Blotting. Ydj1 interacted with Rnr1 equally in Ssa1-4 cells in both unstressed and HU-treated cells (Fig. 5A). Interestingly, both Ydj1-Rnr2 and Ydj1-Rnr4 interaction decreased in cells solely expressing Ssa3 and Ssa4 (Fig. 5B and 5C).

**Figure 5.**
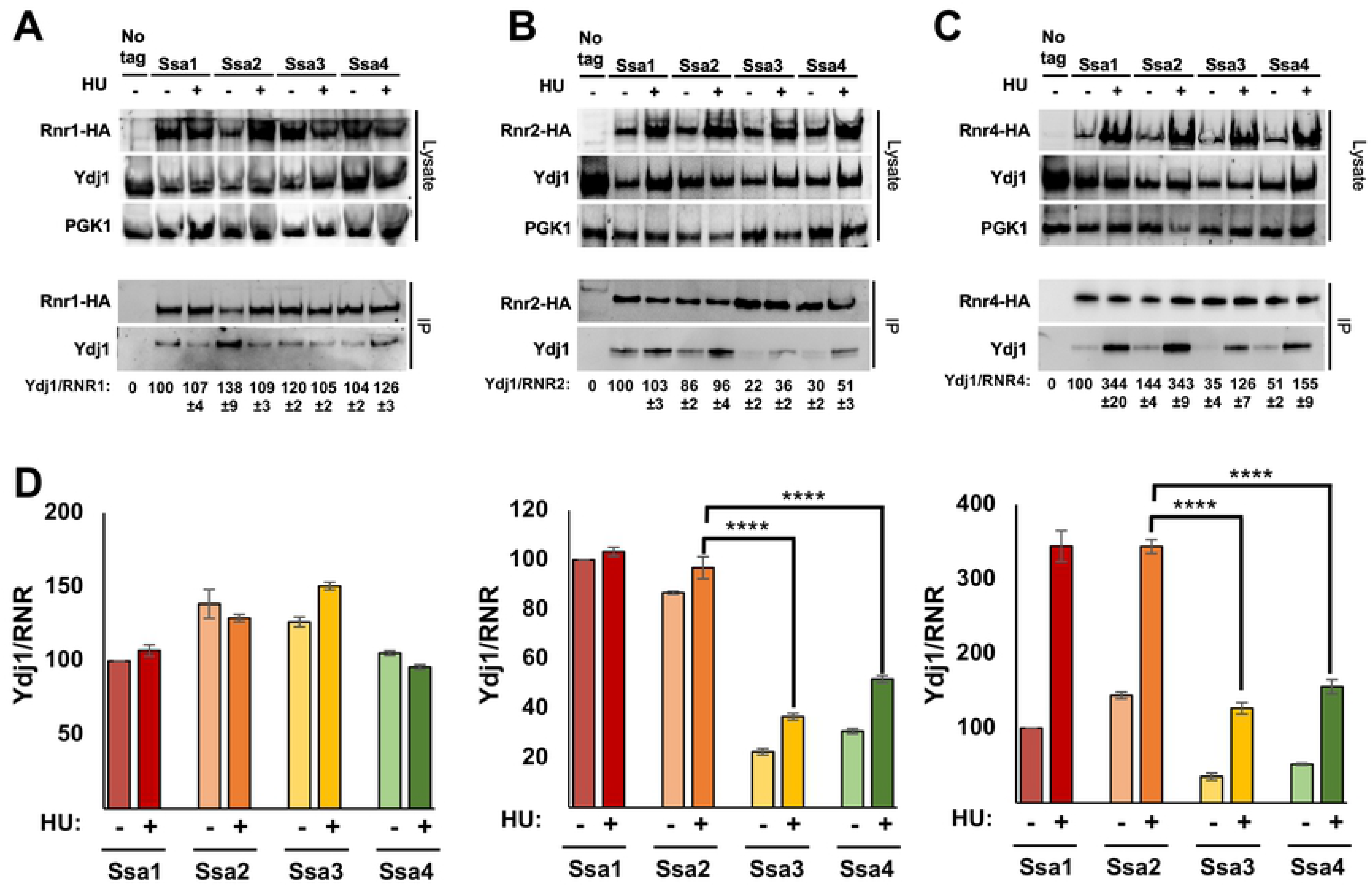
small RNR subunit interaction with Ydj1 is decreased in Ssa3 and Ssa4 cells. (A-C) Cells expressing either Ssa1, 2, 3 or 4 as their sole Ssa and HA-tagged Rnr1/Rnr2/Rnr4 were grown to exponential phase and were either left untreated or were treated with 200 mM HU for 3 hrs. HA-RNR complexes were immunoprecipitated with anti-HA magnetic beads and were subjected to SDS-PAGE and analyzed by immunoblotting with anti-HA antibodies to detect the RNR subunits or anti-Ydj1 antibodies to detect Ydj1. (D) Interaction between RNR and Ydj1 (Ydj1/RNR) was calculated by quantitating bands from three replicate experiment. Each value represents the mean ± SD (n = 3). Statistical significance between samples was calculated ANOVA. (∗∗∗∗ p < 0.01).

### The CTD region of Ssa2 (542-639) is required for full RNR activity

Hsp70 is comprised of 3 major domains, NBD, SBD and CTD, the first of which is connected to the last two via a flexible linker. Ssa2 and Ssa4 share 84% sequence similarity with the SBD being the highest variable region (Fig. 6A and 6B). To understand the origin of the functional difference between the Ssa2 and Ssa4 in regard to RNR function we created Ssa2-Ssa4 chimeras based on [9] and assessed their resistance to media containing HU. The Ssa24 construct consists of amino acids (a.a.) 1-542 of Ssa2 fused to a.a. 543-639 of Ssa4 and vice-versa for the Ssa42 construct. Although yeast expressing the Ssa42 was as resistant to HU as Ssa4 cells, Ssa24 cells phenocopied Ssa4 cells (Fig. 6C). In an effort to determine whether the a.a. region 1-542 of Ssa2 controlled the transcriptional output of the DNA damage response, we compared expression of β-galactosidase driven by a DNA-damage responsive promoter (*RNR3* promoter-lacZ) in HU-treated cells expressing Ssa2, Ssa4, Ssa24 and Ssa42. In correlation with the previous result, Ssa24 cells were unable to fully activate *RNR3* transcription (Fig. 6D). Taken together these results suggest the C-terminal domain of Ssa paralogs is critical in determining the ability to activate RNR activity and thus the cellular response to HU.

**Figure 6.**
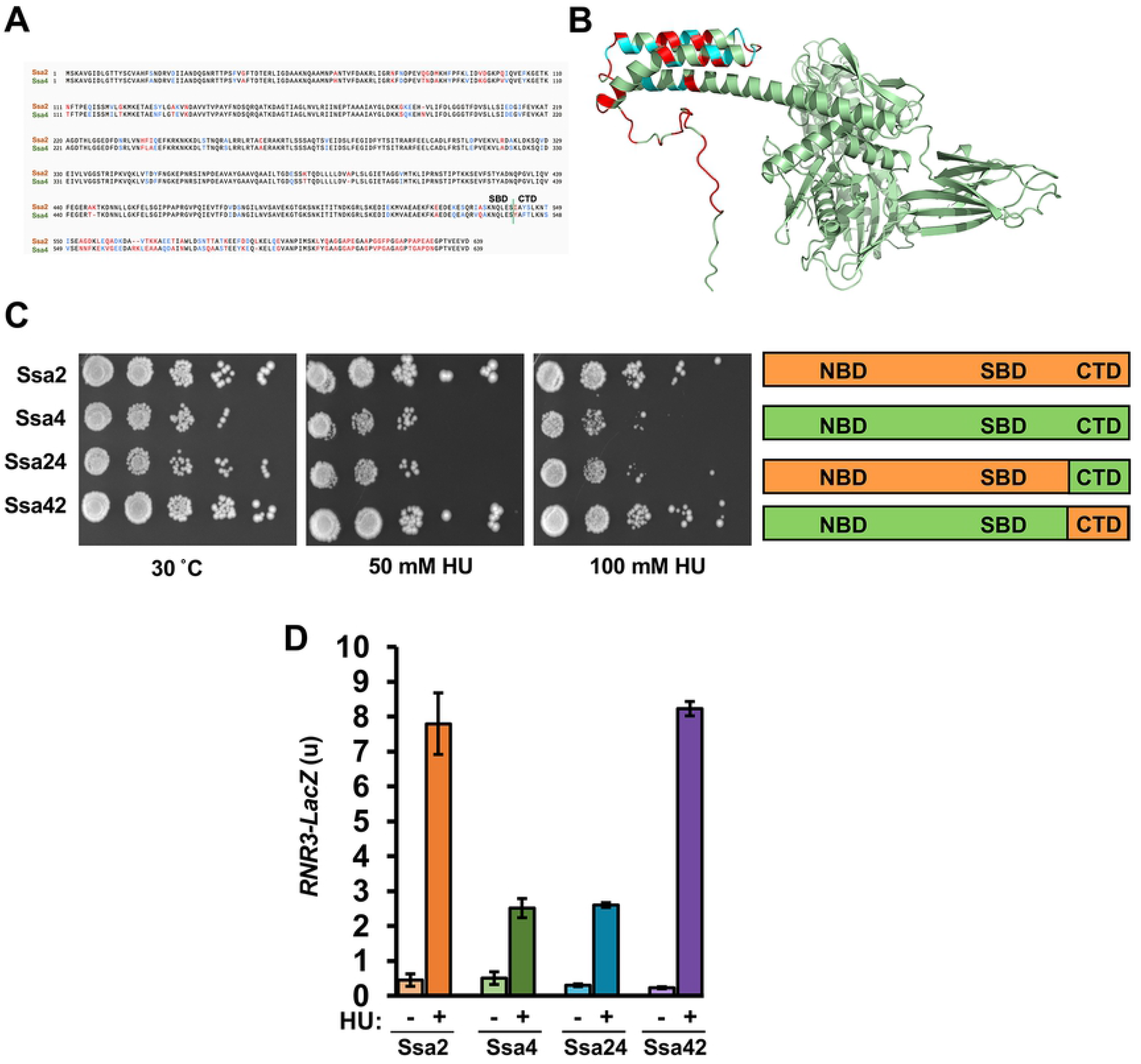
The C-terminus of Ssa2 is required for HU resistance. (A) Sequence alignment between Ssa2 and Ssa4 was created using Clustal Omega. Amino acids are labeled either Black (identical), Blue (similar) or Red (different). (B) Areas of sequence variance between Ssa2 and Ssa4 mapped to the predicted structure for Ssa2. The structure of Ssa2 was modeled via AlphaFold and rendered in PyMol. Residue similarity between Ssa2 and 4 was denoted by color (green, identical; blue similar; red, different). (C) Chimeras of Ssa2 and Ssa4 display altered resistance to hydroxyurea. Cells expressing either Ssa2, 4, 24 or 42 as the sole Ssa were grown overnight to saturation and serial 10-fold dilutions were plated by pin plating from 96-well plates onto YPD alone or YPD containing Hydroxyurea. Plates were imaged after 3 days. (D) DNA-damage response transcription in Ssa2-4 chimeras. An *RNR3-LacZ* reporter plasmid was transformed into the indicated yeast strains. Transformants were grown and subjected to 0 or 200mM HU for 3 hours. β-Galactosidase activity was measured in crude extracts. β-Galactosidase specific activity (in units) [-Gal Sp. Act. (U)] is shown on the y axis. Each value represents the mean and standard deviation (error bar) from three independent transformants; *, P≤0.05**; P≤0.01; ***, P≤0.001 as compared to Ssa2 cell controls.

## Discussion

A fundamental mystery in molecular chaperone research is why cells express so many apparently similar and functionally redundant chaperone paralogs. Historically, it was generally thought that the main differences were in their expression across cells and tissues where the constitutive Hsp70 performed general housekeeping duties and the inducible form protected cells against environmental stress. However, several recent studies have shown that even when chaperone paralogs are expressed in yeast at equivalent levels as the sole cytosolic Hsp70, these cells display dramatically different phenotypes [19–21, 40–42].

A challenge in understanding the differential role of the Ssa paralogs is a lack of verified client proteins. While several Hsp70 interactomes have been published under differing conditions and phosphorylation site mutations, validation of these and their cellular effect is still under investigation [33, 43–47]. Excellent attempts to dissect the roles of Ssa paralogs include the well-established Hsp90 client (Ste11) and a non-yeast client, v-Src in addition to the yeast prions [*URE3*] and [*PSI^+^*] [21]. In our previous studies, we managed to identify Hsp90, Hsp70 and Hsp40 as key regulators of RNR in yeast and humans [35], providing an ideal system in which to further probe SSA isoform-specific differences.

In this study, we observed distinct differences in the ability of yeast expressing single SSA paralogs to survive insults to their genome integrity. Ssa1 and Ssa2-expressing cells were more resistant to HU, 5-FU and H_2_O_2_ than Ssa3 or Ssa4 cells. It is interesting to note that despite the above differences, there appeared to be no difference in the response to MMS between paralogs. This result may reflect the different kinds of DNA damage that these agents inflict on DNA. HU, 5-FU and H_2_O_2_ act primarily by causing single-strand damage and replication stress as opposed to MMS which acts to cause double-strand breaks.

Hsp70 and its corresponding co-chaperone Ydj1 have been shown to play a role in the stabilization of the RNR subunits in both yeast and human cells [34, 35]. In this study, we observed decreased Rnr2 levels and a lack of HU-inducibility of Rnr4 in Ssa3 and Ssa4-expressing cells. Further dissection of this phenomenon revealed that the lowered levels of Rnr2 were primarily due to increased subunit instability as determined by the promoter shut-off experiments. In contrast, the altered levels of Rnr4 in Ssa3/4 cells were a combination of transcriptional and protein stability effects. The lowered transcription of Rnr4 may be a consequence of altered DDR signaling, especially as RNR levels are directly controlled by the activity of Mec1, Tel1, Rad53 and Rad9, the latter of which is an Ssa1/2 client [48]. It is thus possible that in cells lacking Ssa1 and 2 that Rad9 is destabilized leading to an inability to activity DDR and induce Rnr4 expression. While future studies on Ssa paralog interaction with main components of DDR signaling may be informative, our yeast two-hybrid experiments clearly show that Rnr2 and Rnr4 have a binding preference for Ssa1 and Ssa2 compared to their inducible counterparts. Given the amino acid conservation between the four paralogs, such a binding difference is rather striking. Clients of chaperones are processed via their co-chaperones and given that Ydj1 is key for RNR activity, we considered the possibility that paralog-specific binding of RNR subunits may be mediated via Ydj1. Our data in Fig 5 clearly shows this to be the case as Ydj1 interaction with Rnr2 and Rnr4 is decreased in Ssa3/4 cells.

Identifying regions of Hsp70 that determine client specificity remains challenging considering the essential nature of the protein and sequence similarity. Hsp70 is comprised of a nucleotide binding domain (NBD) which is important for co-chaperone binding and ATPase activity, a substrate binding domain (SBD) which is important for client interaction and a C-terminal 10-kDa α-helical subdomain (CTD) that acts as a lid over the binding pocket in the SBD [2]. Out of the four Ssa paralogs, Ssa2 cells are the most resistant to DNA damaging agents including HU, while Ssa4 cells are the most sensitive. In order to further dissect the of the sequence determinants for HU resistance we used Ssa2-Ssa4 chimeras. Interestingly, a chimera consisting of the NBD/SBD of Ssa2 and the CTD of Ssa4 was sensitive to HU, pinpointing the CTD domain of Ssa2 as being key for RNR function and resistance to genome perturbing agents. The highest sequence variation between Ssa2 and Ssa4 occurs within the CTD, specifically the outer-facing region of the “lid” (see Fig. 6). Previous studies have identified this region as being important for the binding of co-chaperone proteins. The VEEVD sequence at the end of Hsp70/Ssa1 is critical interaction with DNAJB-type co-chaperones including Sis1 [49–52]. However, there is also substantial evidence that the CTD is also important for the binding of Ydj1. Loss of the last 8 amino acids of Ssa1 substantially reduces the Ssa1-Ydj1 interaction [53] and a 20-amino acid motif in Ssa1 containing GGAP repeats was recently revealed to be necessary for Ssa1 to bind to Ydj1 and activate both the cell integrity and heat shock responses [54]. These results parallel those seen with mammalian Hsp70 paralogs, where each paralog displays a clear binding preference for certain co-chaperones [39]. Taken together, our data suggest that the sequence variation in the CTD is primarily responsible for differential recruitment and folding of RNR small subunits. It is worth noting that recent studies have uncovered a non-canonical binding site in Hsp70 required for binding and folding of alpha-synuclein [55]. It is possible that this region and the area that determines RNR subunit binding heavily overlap. Studies over the past decade have identified numerous post-translational modifications (PTMs) on chaperones which are collectively known as the “Chaperone Code” [56–58]. This code modifies a variety of chaperone properties including localization, stability, and most importantly client and co-chaperone folding. Several PTMs exist in the C-terminal region of Hsp70 and future studies are required to explore the impact of these PTMs on RNR stability. Although this study was carried out in yeast, Hsp70 and Hsp40 also bind RNR subunits in mammalian cells [34, 35]. We envisage future studies that probe this complex interaction in mammalian cells, possibly in the hope of identifying novel ways to inhibit RNR in cancer, feasible given the role of DNAJA1 in anticancer drug resistance [59]

## Materials and Methods

### Yeast Strains and growth conditions

Yeast cultures were grown in either YPD (1% yeast extract, 2% glucose, 2% peptone) or grown in SD (0.67% yeast nitrogen base without amino acids and carbohydrates, 2% glucose) supplemented with the appropriate nutrients to select for plasmids and tagged genes. Escherichia coli DH5α was used to propagate all plasmids. E. coli cells were cultured in Luria broth medium (1% Bacto tryptone, 0.5% Bacto yeast extract, 1% NaCl) and transformed to ampicillin resistance by standard methods. Hsp70 isoform plasmids pRS315P_SSA2_-SSA1, pRS315P_SSA2_-SSA2, pRS315P_SSA2_-SSA3, pRS315P_SSA2_-SSA4 [60] were transformed into yeast strain *ssa1–4*Δ [61] using PEG/lithium acetate. After restreaking onto media lacking leucine, transformants were streaked again onto media lacking leucine and containing 5-fluoro-orotic acid (5-FOA), resulting in yeast that expressed Hsp70 paralogs as the sole cytoplasmic Hsp70 in the cell.

For tagging the genomic copy of *RNR1*, *RNR2* and *RNR4* with a HA epitope at the carboxy-terminus, the pFA6a-HA-His3MX6 plasmid was used in a manner similar to [34]. A full table of yeast strains and plasmids that were used can be found in Supplemental Table T1. For serial dilutions, cells were grown to mid-log phase, 10-fold serially diluted and then plated onto appropriate media using a 48-pin replica-plating tool. Images of plates were taken after 3 days at 30°C. 200mM HU was used for serial dilutions and to stress yeast cells, a concentration established in Tkach et al. [62].

For IC_50_ calculations, cells were grown to mid-log phase, diluted in a sterile 96 well plate in media containing HU, 5-FU, H_2_O_2_, MMS and were 10-fold serially diluted at indicated concentrations. Cells were continuously shaken for 24 hours at 30°C and the optical density of the reaction was measured at 600nm. The mean and standard deviation from three independent transformants were calculated.

### β-Galactosidase assays

For *RNR3-lacZ* fusion expression experiments, *ssa1-4Δ* yeast cells expressing single Ssa constructs or Ssa2/4 fusions were grown overnight in SD-URA media at 30°C and then re-inoculated at OD600 of 0.2–0.4 and then grown for a further 4 hours. Cells were treated with 150 mM or 200 mM HU for 3 hours and then RNR3-lacZ fusion assays were carried out as described previously in Truman et al. [63]. Briefly, protein was extracted through bead beating and protein was quantitated via Bradford assay. The beta-Galactosidase reaction containing 50 μg of protein extract in 1 ml Z-Buffer (30) was initiated by addition of 200 μl ONPG (4 mg/ml) and incubated at 28°C until the appearance of a pale-yellow color was noted. The reaction was quenched via the addition of 500 μl Na2CO3 (1M) solution. The optical density of the reaction was measured at 420nm. β-Gal activity was calculated using ((OD420 x 1.7)/(0.0045 x protein x reaction time)), where protein is measured in mg, and time is in minutes. The mean and standard deviation from three independent transformants were calculated.

### Galactose promoter shut-off experiments

*Ssa1-4Δ* yeast cells expressing either Ssa1, 2, 3 or 4 as the sole Hsp70 isoform were transformed with either pGAL1-HA-Rnr1, 2 or 4 plasmids were grown to mid-log phase in YP Gal medium (1% yeast extract, 2% galactose, 2% peptone). Transcription of pGAL1-HA-Rnr1, 2 or 4 was shut off by the addition of 2% glucose to cultures. Aliquots of cells were collected at 0 and 4 hours after the addition of glucose. Cell lysates from these samples were analyzed by Western Blotting for stability of RNR subunit (HA antibody) and loading control (PGK1).

### Western blotting

Protein extracts were made as described and 20 μg of protein was separated by 4%– 12% NuPAGE SDS-PAGE (Thermo) [44]. Proteins were detected using the following antibodies; anti-HA tag (Thermo #26183), Anti-FLAG tag (Sigma, #F1365), anti-PGK1 (Thermo # PA5-28612), anti-Ydj1 (StressMarq #SMC-166D). Blots were imaged on a ChemiDoc MP imaging system (Bio-Rad). After treatment with SuperSignal West Pico Chemiluminescent Substrate (GE). Blots were stripped and re-probed with the relevant antibodies using Restore Western Blot Stripping Buffer (Thermo).

### Purification of HA-tagged Rnr1, 2 and 4 from yeast

*Ssa1-4Δ* yeast cells expressing genomically-tagged HA-Rnr1, Rnr2 and Rnr4 were transformed with Ssa1-4 pRS315 plasmids were grown overnight in SD-LEU media, and then reinoculated into a larger culture of selectable media and grown to an OD600 of 0.800. The cells were then either unstressed or stressed with 200 mM HU for four hours.

Cells were harvested and HA-tagged proteins were isolated as follows: Protein was extracted via bead beating in 500 μl binding buffer (50 mM Na-phosphate pH 8.0, 300 mM NaCl, 0.01% Tween-20). 200 μg of protein extract was incubated with 30 μl anti-HA magnetic beads (Sigma) at 4°C overnight. Anti-HA beads were collected by magnet then washed 5 times with 500 μl binding buffer. After the final wash, the buffer was aspirated and beads were incubated with 65 μl Elution buffer (binding buffer supplemented with 10 μg/ml 3X HA peptide (Apex Bio)) for 1 hour at 4° C, then beads were collected via magnet. The supernatant containing purified HA-RNR1, 2, and 4 were transferred to a fresh tube, 25 μl of 5x SDS-PAGE sample buffer was added and the sample was denatured for 5 min at 95° C. 20 μl of sample was analyzed by SDS-PAGE.

### Quantitation of yeast RNR subunit transcription

Quantitation of yeast *RNR* transcription was carried out as in Zhang et al. [64]. Briefly, *Ssa1-4Δ* yeast cells expressing unique Ssas were grown overnight in YPD media at 30°C, re-inoculated at OD600 of 0.2–0.4 and then grown for a further 4 hours. Cells were treated with 200 mM for 2 hours and total RNA was extracted from cells using a GeneJet RNA extraction kit. Total RNA (1 μg) was treated with 10 units of RNase-free DNase I (Thermo) for 30 min at 37°C to remove contaminating DNA. DNAse I activity was stopped by adding 1 μL of 50 mM EDTA and incubating at 65°C for 10 minutes. cDNA synthesis was carried out by iScript reverse transcriptase (BioRad) on aliquots of 1 μg RNA. The single-stranded cDNA products were used in qPCR on an ABI Fast 2000 real-time PCR detection system based on SYBR Green fluorescence. Sequences of oligo pairs (same as used in [64]) are listed in S1 Table. Signals of *RNR1*, *RNR2* and *RNR4* were normalized against that of *ACT1* in each strain and the resulting ratios in WT cells were defined as onefold.

### Yeast Two Hybrid Analysis

*Ssa1-4Δ* cells expressing unique Ssas were transformed with the appropriate GAL4 AD and BD fusion proteins. Interaction between Ssa paralogs and RNR subunits was measured via β-galactosidase assays as in [63].

## Abbreviations

RNR: Ribonucleotide reductase
DDR: DNA damage response
IP: immunoprecipitation
dNTP: deoxyribonucleotide
HSP: Heat Shock Protein
ER: Endoplasmic Reticulum

## Acknowledgements

We would like to thank Deepak Sharma and Daniel Masison for their reagents and advice on this work.

